# Single-cell dissection of a collective behaviour in honeybees

**DOI:** 10.1101/2022.03.29.486106

**Authors:** Ian M. Traniello, Syed Abbas Bukhari, Payam Dibaeinia, Guillermo Serrano, Arian Avalos, Amy Cash Ahmed, Alison L. Sankey, Mikel Hernaez, Saurabh Sinha, Sihai Dave Zhao, Julian Catchen, Gene E. Robinson

**Affiliations:** Neuroscience Program, University of Illinois at Urbana-Champaign; Urbana, IL (UIUC); Carl R Woese Institute for Genomic Biology, UIUC; Department of Computer Science, UIUC; Computational Biology Program, CIMA University of Navarra; Pamplona, Spain; United States Department of Agriculture – Agricultural Research Services, Honey Bee Breeding, Genetics and Physiology Research Laboratory; Baton Rouge, LA; Department of Statistics, UIUC; Department of Evolution, Ecology and Behavior, UIUC; Department of Entomology, UIUC

## Abstract

Understanding how genotypic variation results in phenotypic variation, a major challenge in biology, is especially difficult for collective behaviour because collective group phenotypes arise from complex interactions between group members^1^. Honeybees aggressively defend their colony from attacks with highly integrated collective behaviour in which different groups of bees play specific roles, giving rise to distinct colony-level differences in aggression. A previous genome-wide association study of a population of Africanized honeybees (*Apis mellifera scutellata*) from Puerto Rico that recently evolved decreased aggression identified hundreds of genes with single nucleotide polymorphisms (SNPs) that associated with colony-level variation in aggression^2^. Many of these SNPs also showed strong signals of selection for decreased aggression^2,3^, but their influence on brain function was unknown. Using brain single-cell (sc) transcriptomics and sc gene regulatory network analysis, we show here that variants of these genes give rise to genetic differences in transcription factor-target gene relationships. These differences involved the activity of several TFs, some that have been previously associated with aggression, like *single stranded-binding protein c31A*, and some that have been associated with tissue morphogenesis but not behaviour, like *apontic*. The activity of these and other TFs was located in specific brain cell populations related to olfaction and vision, the two sensory modalities that bees use in colony defence. They also implicate metabolism of serotonin, a neurochemical already known to influence honeybee aggression, but not from a genetic perspective. Surprisingly, genetic differences were more pronounced in the brains of forager bees than in similarly aged but more aggressive soldier bees, pointing to an evolutionary change in division of labour for colony defence. Our results demonstrate how group genetics can shape a collective phenotype by modulating individual brain gene regulatory network architecture.

## Main

In honeybee societies, individual workers are integrated into collective efforts via division of labour^4,5^. One such example is colony defence following territorial intrusion, in which highly aggressive soldier bees mount a rapid stinging attack while similarly aged nestmates ignore the disturbance and instead forage for nectar and pollen. Colony-level variation in aggression is well known in honeybees, and both inherited and environmental determinants have been identified^2^. This is consistent with Darwin’s assertion that natural selection acts at the colony level in the insect societies^6,7^, but it is not known at the genomic level how colony-level selection shapes the phenotypes of individual colony members. We used specific variants of genes that associated with variation in colony aggression (“colony aggression genes”) to explore this problem.

We tested the prediction that variation in collective behaviour manifests in the neurogenomic architecture of high-aggression soldiers and low-aggression foragers by using the recent evolution of decreased aggression in the otherwise highly aggressive Africanized honeybee (AHB)^3^. We compared whole-brain transcriptomic profiles of 81 soldiers and 82 foragers from nine Puerto Rican honeybee colonies, using the same individuals previously subjected to a whole-genome analysis^2^. We coupled this analysis with scRNA-Sequencing to profile the transcriptomes of ∼24,000 cells from 40 soldier and forager brains from four additional high- and low-aggression colonies in Urbana, Illinois (fig. S1; fig. S2A-B; Data S1). This allowed us to detect differentially expressed genes (DEGs) between soldiers and foragers, both in bulk brain samples and resolved to the level of specific brain cell populations. Neither DEG list overlapped significantly with the colony aggression genes (Fig. 1A; fig. S2C; Data S2-S3).

**Fig. 1.**
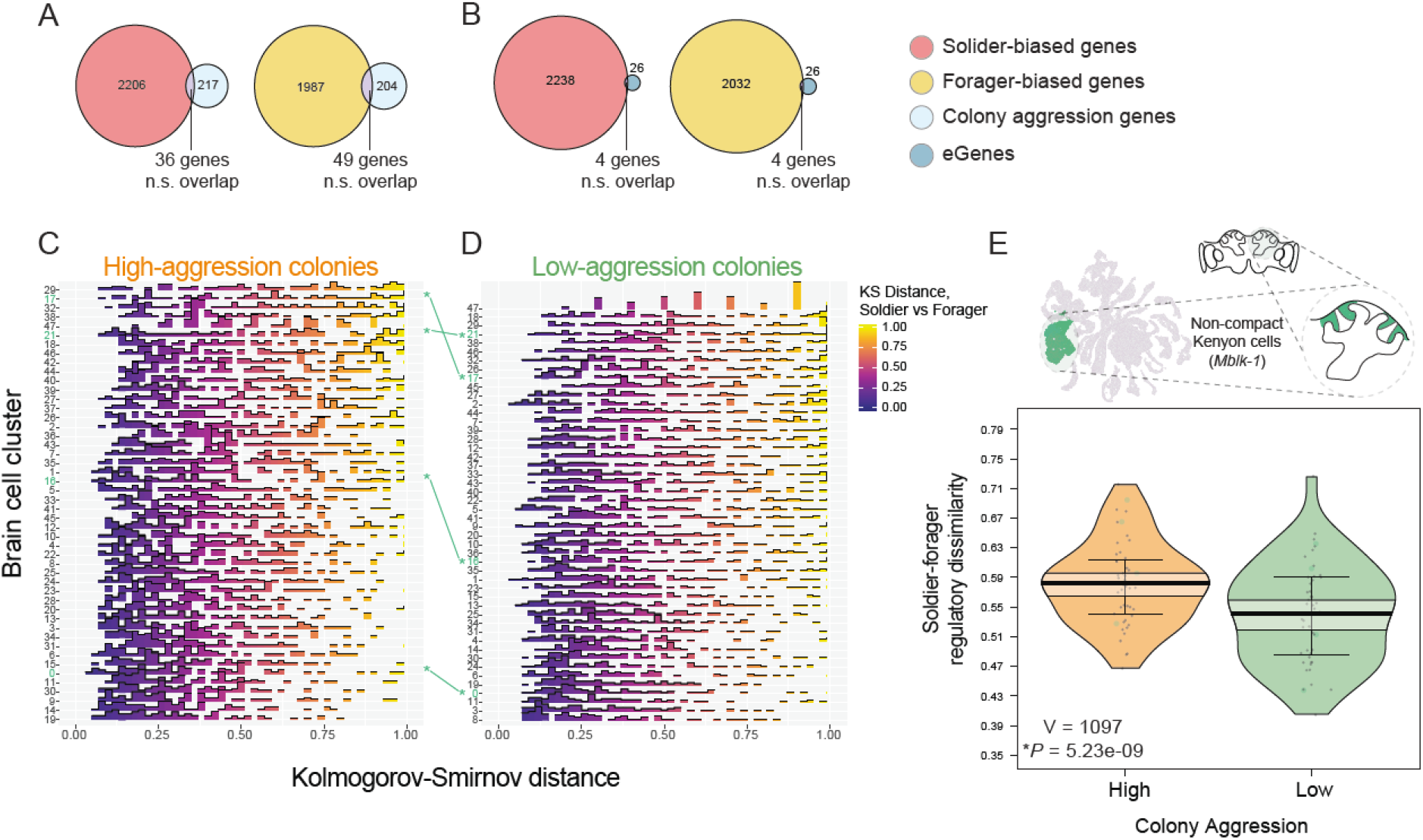
Dissimilarity of brain gene regulatory network (GRN) architecture between soldiers and foragers is greater in high-aggression honeybee colonies. (A) No significant enrichment of genes containing colony aggression-associated SNPs (“colony aggression genes”) or (B) of eGenes in soldier or forager-biased differentially expressed genes (proportional Venn diagrams; hypergeometric test, *P* > 0.10). (C-D) By contrast, genetic differences in colony aggression are manifest in brain single-cell GRNs. Distribution of Kolmogorov-Smirnov (KS) regulatory dissimilarity (RD) distances, representing variation in brain GRN architecture between high-(soldiers) and low-aggression (foragers) individuals, showing greater RD between soldiers and foragers from high-compared to low-aggression colonies. Each row represents one of 48 cell clusters from four soldier and four forager scRNA-Seq samples from two high-and two low-aggression colonies. Numbers highlighted in green with asterisks: lines show relative position of specific brain cell clusters as a function of RD in high-and low-aggression colonies. (E) *Above*. UMAP plot and brain diagram provide heuristic example of brain cell populations with greater soldier-forager RD associated with greater colony aggression, highlighting the non-compact Kenyon cells (ncKCs, identified via expression of *Mblk-1*). *Below*. Aggregated data from panels C and D showing significantly greater RD in bees from high-versus low-aggression colonies (Wilcoxon signed-rank exact test). Solid line in center of plots = mean; shaded bands = 95% confidence interval; whiskers = 1^st^ and 3^rd^ quartiles; and raw data shown as points with surrounding smoothed density curve. Each point is derived from a row in corresponding panels C and D. Larger green points represent cell clusters belonging to the ncKCs.

We next used expression quantitative trait loci (eQTL) mapping to search for specific genes linking variation in group genetics to variation in individual brain function^8^. We first identified thousands of eQTL-genes (eGenes) containing *cis*-located SNPs (eSNPs) in soldiers and foragers, and filtered this list to 30 eGenes by requiring them to have eSNPs that were also SNPs in the colony aggression genes^2^ (Data S4). Like for colony aggression genes, we did not find a significant overlap with the soldier-forager DEGs at bulk or sc resolution (Fig. 1B; fig S2C). We therefore asked whether genetic differences in colony aggression manifest at the level of gene regulatory networks (GRNs).

Recent theory predicts that the genetic architecture of polygenic traits, including most behaviours, is best understood at the level of the GRN, which models regulatory relationships between transcription factors (TFs) and their target genes (TGs)^9,10^. This prediction is consistent with an emerging appreciation that behaviourally related GRNs are an important level of organisation in the brain^11^. Developmental GRNs have proven to be useful in the study of evolutionary changes in morphological traits^12^, and variation in DNA sequence has been associated with subtle changes in GRN activity underlying disease states^13,14^. However, GRN analysis has not yet been employed to study how changes in the brain affect collective behaviour.

We modeled GRNs from the sc data and determined the regulatory dissimilarity score, a measure of network difference that integrates across all cell-type-specific TF-TG relationships^15^. There was greater GRN dissimilarity between soldiers and foragers from more aggressive colonies relative to less aggressive colonies (Fig. 1C-E; fig. S3). This finding indicates that the collective environment is related to brain gene regulatory architecture for individual colony members, an example of an indirect genetic effect^1^ acting on the brain. These genetic differences in regulatory architecture were prominent in two cell populations of note: the non-compact Kenyon Cells (KC) of the mushroom bodies (MB) and astrocyte-like glia. The former have been associated with honeybee aggression via *in situ* hybridization^16^ and the latter have been associated with behavioural plasticity in honeybees^17^ and ants^18^, but never before from a genetic perspective.

Having established a relationship between genetic differences in colony aggression and differences in GRN structure, we tested for an influence of the 30 eGenes on the brain gene regulatory architecture underlying the soldier and forager behavioural states. We developed a new computational method, similar to ref. ^19^ and tested the prediction that if eGenes act in concert they will form tightly connected regulatory subnetworks, measured by the strength and degree of TF-TG relationships distributed across different brain cell populations. We analyzed GRN subnetworks composed of either the 30 eGenes as TGs or, as a negative control, 15 random permutations of equally sized subnetworks using eGenes with eSNPs that did not overlap with SNPs in the colony aggression genes^2^. As predicted, the eGene subnetworks were highly similar to each other in terms of TF-TG relationships, more so than the random subnetworks, which did not show any cohesive signal (Fig. 2A-B; fig. S4). This finding reveals a mechanism by which group genetics can shape brain and behaviour by influencing the functional integration of GRNs.

**Fig. 2.**
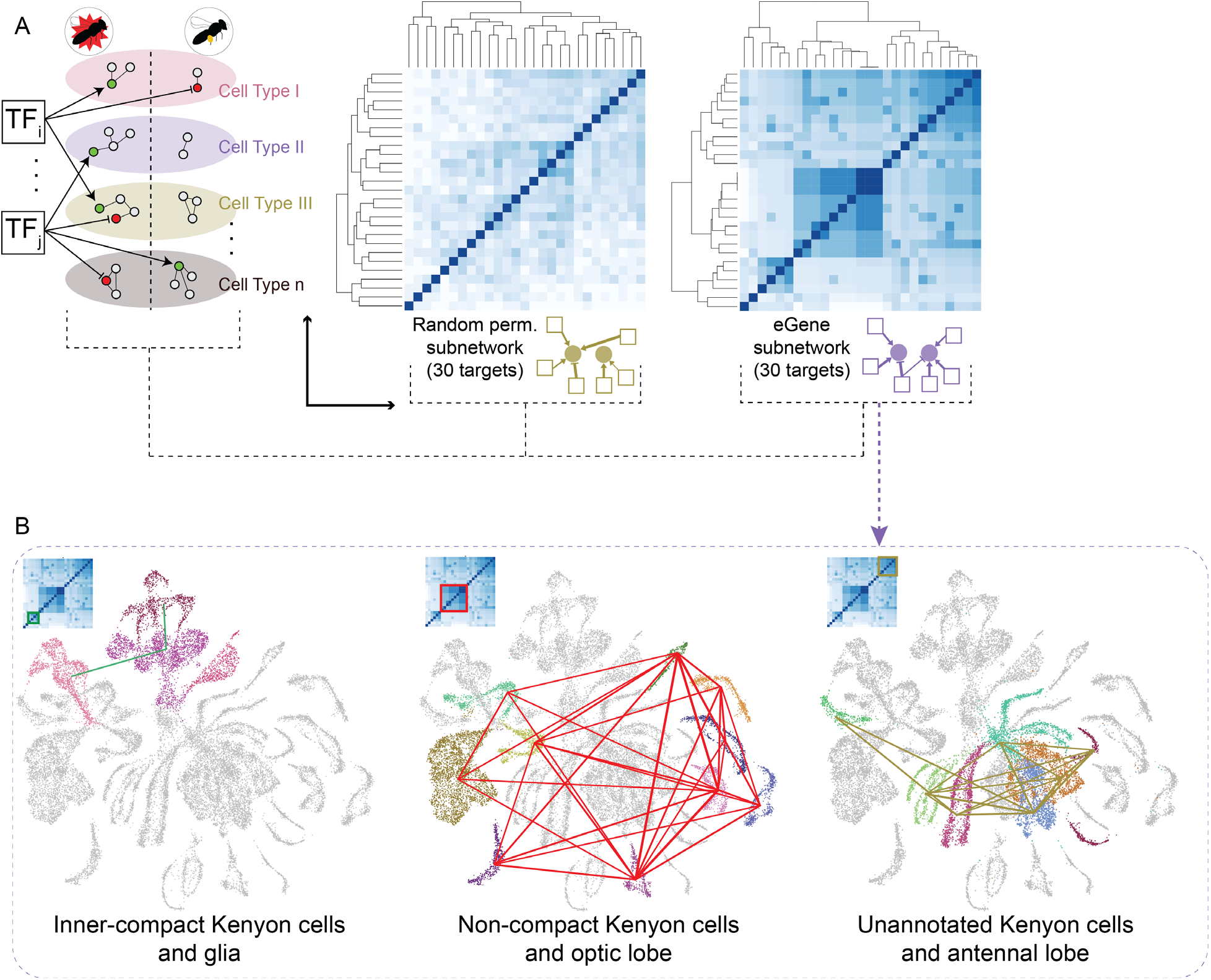
Evolutionary changes in honeybee colony aggression are associated with changes in brain cell gene regulatory network (GRN) architecture between soldiers and foragers. (A) Subnetworks composed of either eGenes or random subsets of genes (negative control) as target genes (TGs) were extracted from an inferred scGRN (Methods). Heatmaps comparing these subnetworks across cell clusters reveal negligible similarities for the control subnetworks (representative example from permutation analysis shown here) but highly similar architectures for the eGene subnetworks. Darker spots represent more similar sets of Transcription Factor-TG relationships across cell clusters, as determined by Jaccard similarity index. (B) Results from (A) superimposed on UMAP plots in which line thickness is proportional to the similarity of subnetwork architecture between two connected cell clusters. For visualization purposes, only edges with similarity ≥ 0.60 are shown.

The eGene subnetworks featured four highly connected TFs: *Single stranded-binding protein c31A* (*Ssb-c31a*), *apontic* (*apt*), *ecdysoneless* (*ecd*), and *CG31460* (Fig. 3A; Data S5). *Ssb-c31a*, which has been associated with population-level variation in honeybee aggression^20^, was most prominent in astrocyte-like glia, already identified above, and perineurial glia. The TF *apt*, which has been associated with tissue morphogenesis in *D. melanogaster*^21^ but has not yet been implicated in social behaviour, was most prominent in the inner-compact KC, a small population of MB neurons flanking the above-mentioned non-compact KC which also have been implicated in honeybee aggression^16^, and in the antennodorsal projection neurons of the antennal lobe, which communicate olfactory sensory information to the KC. Overall, there were 15 TF-TG relationships in the eGene subnetworks that showed strong soldier-forager differences, including *Ssb*-*c31a* and *apt*, which also showed the strongest expression differences between soldiers and foragers (Data S6).

**Figure 3.**
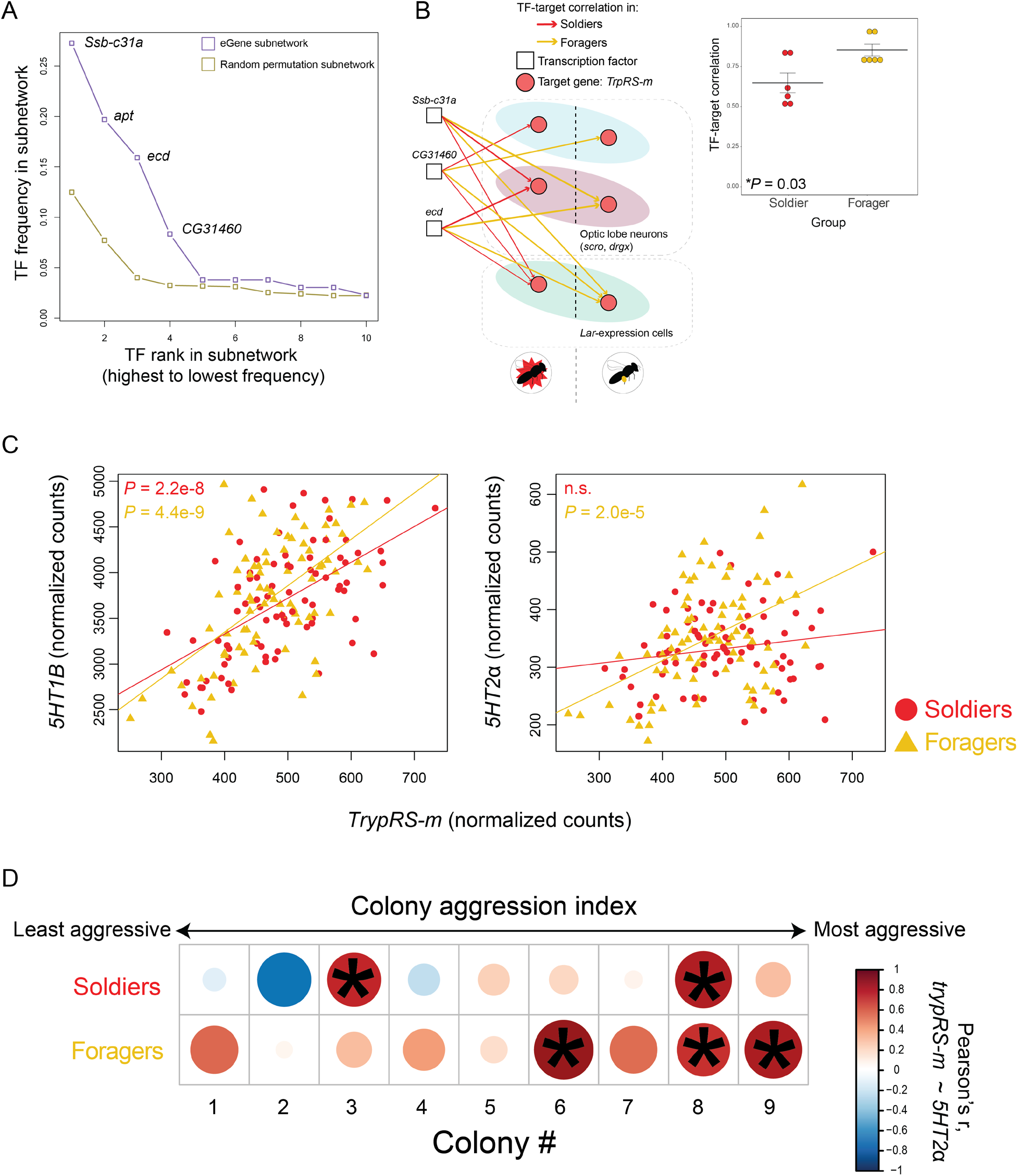
Serotonin metabolism as a representative example of the relationship between colony genetics, regulatory plasticity, and individual behavioural variation. (A) Analysis of TF centrality, as measured by frequency of a given TF, revealed four TFs overrepresented in eGene subnetworks relative to control subnetworks. (B) Strong regulatory plasticity of the TG *TrpRS-m* between soldiers in foragers in three cell clusters identified as part of the honeybee optic lobe; line thickness corresponds to strength of TF-TG relationship. Inset shows that the strengths of these regulatory relationships are significantly higher in foragers compared to soldiers (Wilcoxon). Sample sizes as in Fig. 1. (C) Normalized expression of the serotonin receptor genes *5HT1B* (left) and *5HT2α* (right), both of which were differentially expressed in foragers compared to soldiers, were correlated with expression of the tryptophan metabolism gene *TrypRS-m* in behaviour-specific context: while a significant correlation was detected in both soldiers and foragers for *5HT1B*, it was only detected in foragers for *5HT2α*. (D). When broken down by colony, the correlation between *5HT2α* and *TrypRS-m* was stronger in foragers than soldiers for the most aggressive colonies. Circle size corresponds to absolute value of correlation, and only significant (Pearson’s correlation, \**P* < 0.10) positive associations are shown.

Also identified in the eGene subnetwork analysis was the TG *tryptophanyl-tRNA synthetase mitochondrial* (*TrpRS-m*), the expression of which has been correlated with the expression of genes associated with availability and reuptake of tryptophan^22^, the precursor to the neurochemical serotonin. Serotonin is involved in the regulation of aggression in diverse species of vertebrates and invertebrates^23^, including honeybees^24^. The strongest soldier-forager differences in TF-TG relationships for *TrpRS-m* were in cell types that contain optic lobe markers^25^ and a population of cells primarily expressing *leukocyte-antigen-related-like* (*Lar*), a receptor tyrosine phosphatase required for neuronal development of the visual system in *Drosophila melanogaster*^26^. eGene subnetwork analyses indicate that inherited changes in honeybee aggression are based, in part, on changes in the structure and function of physiological pathways that respond to or process aggression-triggering olfactory and visual stimuli.

The evolution of decreased colony aggression in the Puerto Rican population involved a soft selective sweep^3^, so it is likely that colony-level selection had multiple influences on the brains of colony members. Surprisingly, selection apparently affected the brains of foragers more than soldiers, based on the following four results: 1) genes with expression levels that correlated with colony aggression were significantly enriched for eGenes in foragers, but not soldiers (fig. S5; Data S7); 2) the regulation of *TrpRS-m* expression was significantly stronger in foragers compared to soldiers (Fig. 3B); 3) serotonin-related genes were significantly upregulated in the brains of foragers compared to soldiers (fig. S1A; Data S1); and 4) expression of the serotonin receptor *5HT2α* more strongly correlated with expression of *TrypRS-m* in foragers than in soldiers, and this correlation was most pronounced in foragers from highly aggressive colonies (fig. 3C-D). These results make sense in light of how division of labour works in honeybee colonies. Unlike in ant and termite societies, there are no morphological differences among honeybee workers^27^, so variation in individual aggression is in degree and not kind. Honeybee colony aggression and foraging intensity are inversely correlated because foragers join in defending their colony if triggering stimuli exceed their threshold of response^28^. These results suggest that colony genetics affects the propensity of foragers to participate in defending their colony, in addition to affecting the propensity of soldiers to initiate and sustain the group response.

How the behaviour of individuals is integrated into a smoothly functioning group is one of the central issues in the study of collective behaviour^27^. Our study shows how genomics can be used to analyze a collective behaviour that has undergone evolutionary change by using genotype-phenotype mapping that focuses on GRN architecture. This genotype-phenotype mapping was not possible with analyses based only on gene expression. The approach used here, which uses scGRNs to integrate population genetics and systems biology, provides tools to better understand the evolutionary tuning of a collective behaviour.

## Acknowledgements

We thank A. Hernandez and C. Wright at the Roy J. Carver Biotechnology Center (UIUC) for sequencing services, C. Desplan for useful discussion, and M.B. Sokolowski, members of the Gene Networks in Neural and Developmental Plasticity theme (IGB), and the Robinson lab for valuable comments and feedback on the manuscript.

## Author contributions

Conceptualization: IMT, AA, GER

Methodology: IMT, SAB, PD, GS, ACA

Investigation: IMT, ACA, AS

Visualization: IMT, PD, GS

Funding acquisition: GER

Project administration: GER

Supervision: MH, SDZ, SS, JC, GER

Writing – original draft: IMT, GER

Writing – review & editing: IMT, SAB, PD, GS, AA, ACA, MH, SS, SDZ, JC, GER

## Data availability

Whole-tissue and single-cell RNA-Sequencing data will be made accessible following acceptance for publication.

## Supplementary Materials for

### Other Supplementary Materials for this manuscript include the following

Data S1 to S7

Data S1: Cluster-defining differentially expressed genes and cell cluster annotations

Data S2: Soldier-and forager-biased genes differentially expressed in pairwise comparison Data S3: Cluster-specific differentially expressed genes (soldiers compared to foragers) Data S4: Results from eQTL analysis

Data S5: TF-target interactions in eGene GRN subnetwork

Data S6: Fifteen TF-target interactions that strongly covary with behavioural state

Data S7: Genes with expression levels that correlate with colony aggression (in all individuals and resolved to soldiers or foragers)

## Materials and Methods

### “Gentle” African honeybee (gAHB) soldier and forager collections

Collection strategies for gAHB soldiers and foragers used for the previously published whole genome sequences are described in ref. ^2^. Briefly, nine colonies from eight locations were sampled across Puerto Rico in April 2017, and all colonies were maintained in standard Langstroth hives containing a single hive box with 16,000 – 20,000 bees and one naturally mated queen. Colonies were transported to the Gurabo Agricultural Field Station at the University of Puerto Rico, spaced > 1 m apart, and allowed to rest undisturbed for 2 weeks to acclimate to the new conditions. Queen presence was determined before and after the phenotyping.

A previously described assay for collecting soldiers and foragers was deployed for individual phenotyping^29,30^. A string of nine 5 × 5 cm suede leather patches was positioned 21 cm from the colony entrance, and the colony was rhythmically struck with a cinder block for 40 times over 2 min. Bees that rushed out to sting the patches over the next 2 min (“soldiers”) were collected and flash-frozen in liquid nitrogen, after which the leather patches were removed and placed in a plastic bag for further analysis. The hive entrance was then restricted and bees that left for the next 30 s were dusted with talcum powder. Next, the hive entrance was opened, and the colony was left for 30 min, after which the entrance was sealed again and talcum powder-dusted bees returning to the colony with a pollen load (“foragers”) were collected and flash-frozen. This strategy allowed for the selective collection of foragers that were in the colony during the disturbance but did not respond to it and instead initiated foraging. Frozen samples were stored at the University of Puerto Rico, Rio Piedras, before being transferred to the University of Illinois at Urbana-Champaign (UIUC) for DNA^2^ and RNA processing, the latter of which is presented here.

Stings in the black suede leather patches were counted to measure the intensity of each colony’s response. Two weeks later, colonies were scored again with an established behavioural ranking system^29,31^, which assigns a colony rank score on a scale from one to four based on observations of bees (1) running on the comb, (2) hanging from the comb, (3) flying around the hive, and (4) stinging investigators. Behaviours “1” and “2” are markers of general activity, “3” is associated with stronger arousal, and “4” is a measure of colony aggression during a lower level of disturbance than previously administered with the colony disturbance assay. Colonies were assigned a cumulative rank score from 1-9 following criteria from each assay.

A multidimensional scaling (MDS) analysis was then applied to generate a unified colony phenotype from the combined colony measures. The first dimension generated by the MDS approach was highly correlated with colony rank across the two measures, and we used either this dimensional vector or number of stings in the leather patches as a surrogate for colony-level aggressive phenotype, as noted. Full experimental details including colony scores, ranking, and MDS dimensional vectors are available in ref. ^2^.

### Bulk RNA-Sequencing on gAHB samples

We used gAHB soldier and forager samples from Puerto Rico that had been previously collected for whole-genome sequencing^2^, as described above. Heads from individual bees were removed and freeze-dried for 52 min at -80°C before the brain was dissected in a dry ice/ethanol bath. Total brain RNA was then extracted using the RNeasy Mini Kit (QIAGEN, Hilden, Germany) and eluted in 30 ul of RNase-free water. Quality and quantity of RNA was assessed on a NanoDrop 1000 (Thermo Fisher, Waltham, MA), Qubit 2.0 fluorometer (Thermo Fisher), and Bioanalyzer (Agilent Technologies, Santa Clara, CA). RNA-Sequencing (RNA-Seq) libraries were prepared on two plates with the TruSeq Stranded mRNAseq Sample Prep Kit, and libraries were quantitated via qPCR. We performed sequencing on one lane of a NovaSeq 6000 with the NovaSeq S4 reagent kit for 151 cycles from each end of the fragments. Reads were 150 nucleotides in length. We demultiplexed FastQ files with the bcl2fastq Conversion Software v2.20 (all Illumina unless otherwise stated).

### European honeybee (EHB) soldier and forager collections

For collections of EHB soldiers and foragers in Urbana, IL, we used the same approach outlined in ref. ^2^, as detailed above, which describes the gAHB samples referenced above and the bulk brain RNA-Seq analysis used in this study. All EHB samples were used for single-cell RNA-Seq (scRNA-Seq).

All colonies were maintained according to standard beekeeping practices at the Bee Research Facility at UIUC. We relocated four standard Langstroth hives containing a colony of ∼30,000 workers headed by a naturally mated queen to a common yard to control for environmental factors like floral resource access and exposure to ambient conditions. Hives were equal in size, containing two stacked hive boxes with identical amounts of honeycomb and were left undisturbed for at least 10 days after relocation and prior to phenotyping. We collected soldiers and foragers from one colony per day across four successive days in August 2019 (n = 4 soldier and 4 forager samples). On each collection day, we began at 10:00AM by performing the colony disturbance / stinging assay as described above. However, instead of being flash-frozen, samples were rapidly transported indoors and processed for scRNA-Seq.

### Single-cell RNA-Sequencing (scRNA-Seq)

Soldier and forager brains were dissected in ice-cold Dulbecco’s PBS (DPBS, cat. no. #14190144, Thermo Fisher Scientific) in a sterile, RNase-free space. We then followed the “short dissociation protocol” from^25^, which allowed us to go from sample collection in the field to cells in suspension within minutes. Briefly, we cleaned away glands, trachea, and any connective tissue and cut each freshly dissected brain into six pieces, leaving two optic lobes and the central brain divided into four quadrants. We were careful to only cut through neuropil, thus leaving neurons and neuron-dense regions intact. Brain pieces were transferred to a sterile centrifuge tube containing 500 ul of DPBS on ice. After this was repeated for five brains per soldier or forager sample, we allowed all tissue to sink to the bottom of the tube and replaced DPBS with 200 ul of a diluted Trypsin-EDTA solution (0.05%, cat. no. 25300054, Invitrogen, Carlsbad, CA) and transferred the centrifuge tube to a thermomixer. Samples were mixed at 1000 rpm and 25°C for 15 min with gentle manual trituration with an ordinary pipette tip every 5 min. We then removed residual tissue from the bottom of the tube with a wide-bore pipette tip and centrifuged cells at 400 rcf and 4°C. We removed the supernatant and washed the cells twice in 1 ml ice-cold DPS with a centrifugation step in between washes. After the second wash, we performed a final centrifugation step and resuspended cells in 400 ul DPBS with 0.04% UltraPure™ Bovine Serum Albumin (AM2616, Thermo Fisher Scientific). We then filtered cells with a 40 Flowmi™ Tip Strainer (H136800040, Bel-Art, Wayne, NJ). Cell suspensions were stored on ice and immediately transported to the Roy J. Carver Biotechnology Center at UIUC, where they were subjected to a total cell count and % viability assay with AOPI staining on a Nexcelom Cellometer Spectrum Image Cytometry System (Nexcelom Biosciences, Lawrence, MA USA). Viability was consistently > 97% for each sample.

Libraries were constructed with the Chromium Single-Cell V3 kits (10x Genomics, Pleasanton, CA) and the library pool was amplified via qPCR and loaded on one lane of a NovaSeq 6000 S4 flowcell (Illumina, San Diego, CA), where it was sequenced for 28 nt from one end and 151 nt from the other end. FastQ files were generated and demultiplexed with the CellRanger v3.0.2 “mkfastq” command. The pool of libraries produced over 5 billion reads, and excellent quality was observed for each sample via fastQC.

Within a few days after collections, we inspected all four colonies and confirmed the presence of a laying queen, meaning each colony was queenright at the time of sample collection.

## Bioinformatics

### Bulk RNA-Seq

We removed 12 samples from analysis due to contamination with a common honeybee virus^17,32^ and two samples with an abnormally low number of sequenced reads, leaving us with a final sample size of 163 individuals (81 soldiers and 82 foragers) collected from nine colonies. We identified and removed a batch effect from library preparation plate using Combat-Seq^33^ before performing differential gene expression analysis with edgeR^34^. After filtering to remove transcripts that had fewer than one count-per-million in at least the number of samples equal to the smallest group size, we performed TMM normalization on a gene list universe of 9185 genes for pairwise soldier versus forager comparisons and 9827 for an analysis of genes correlated with colony aggression in soldiers or foragers, the latter being larger on account of smaller group sizes and therefore a more relaxed filtering. We used a generalized linear model (GLM) to compare gene expression between soldiers and foragers using colony as a blocking factor. We next independently analyzed soldiers and foragers to identify genes positively or negatively associated with aggression by regressing individual-level gene expression on each individual’s colony aggression value, derived from the MDS dimensional vector from^2^, as a continuous predictor of gene expression. To correct for multiple tests, we performed the Benjamini-Hochberg (BH) correction ^35^ and filtered differentially expressed gene (DEG) lists based on a false discovery rate (FDR) threshold of 0.05. For Gene Ontology (GO) enrichment analysis^36^, we first converted soldier-or forager-biased honeybee genes to *Drosophila melanogaster* orthologues using a one-to-one reciprocal best hit BLAST (RBH-BLAST). Enrichment of terms associated with Biological Processes was performed with GOrilla ^37^ and visualized with REVIGO^38^.

Gene list overlap analysis was performed via hypergeometric tests with BH-FDR correction for multiple tests using the GeneOverlap package^39^. This approach was also used to compare bulk and single-cell data. We directly compared gene lists associated with behavioural state, colony aggression level, or cell cluster (described below) with a list of 253 genes that contained aggression-associated single-nucleotide polymorphisms (SNPs); 60 of these genes also showed significant signatures of selection, meaning their frequency was found to be increased in the gAHB population. We refer to these 253 genes as “colony aggression genes,” and the complete list can be found in Dataset S1 of ref. ^2^.

### Expression quantitative trait loci (eQTL) mapping

To test for an association between genotype and gene expression, we accessed the genotype calls made on the same individuals as used in this study from ref. ^2^. eQTL-SNPs (eSNPs) with variance greater than zero were retained for the eQTL analysis, and a distance of 10kb was used to define local SNP-gene pairs. A linear model was fitted in Matrix eQTL^40^ to scan for genome-wide *cis*-eQTLs for 9185 genes and 3309291 SNPs separately for soldiers and foragers. Individual relatedness and different colonies were blocked in the linear model and an FDR cutoff of 0.05 was used to select the *cis*-eQTLs.

This analysis identified 3935 and 3473 eQTL-genes in soldiers and foragers, respectively. To identify variants most likely to be associated with both group-and individual-level aggression, we short-listed the eQTL-genes by intersecting eQTL variants in both soldiers and foragers with the variants associated with colony aggression in ref. ^2^. This resulted in 30 eGenes which were used for downstream analysis. We tested for overlap enrichment in the bulk and sc transcriptomics experiments, and the 30 eGenes were also used as TGs in the gene regulatory network analyses described below.

### Single-cell RNA-Sequencing

scRNA-Seq data were aligned, filtered, and quantitated using the CellRanger “count” command. To assess cell quality, we considered the expression of both mtDNA as well as mitochondrially localized proteins, as previous studies have suggested both may be associated with cell death or dysfunction^41–43^. We detected exceptionally low levels of each (combined averages: ∼2 ± 1%) which, when taken with our viability analysis, indicates that the vast majority of sequenced cells were live and yielded high quality data.

Downstream analyses were performed with Seurat^44^.We required each gene to be detected in at least 3 cells and each cell to contain at least 200 genes. Data were log normalized via the *NormalizeData* command with default settings, and the mean-variance relationship was controlled for by running *FindVariableFeatures* with default parameters. To correct for batch effects based on day of sequencing, we used the “IntegrateData” function^44^ which simultaneously controlled for variation due to colony of origin, as each of the four sequencing days was dedicated to a different colony. We then constructed a Shared Nearest Neighbor graph and applied the Louvain community detection algorithm to cluster cells before visualizing data with Uniform Manifold Approximation and Projection for Dimension Reduction (UMAP). UMAP plots were generated using the top 50 dimensions from a principal component analysis and “min.dist” set to 0.75 to evenly distribute cells for visualization. Clustering resolution was set to the default value of 0.8. These parameters were selected based on previous scRNA-Seq data in *D. melanogaster*^25^, jumping ants^18^, and a previous honeybee study^17^, with consideration for the high degree of cellular homogeneity in insect whole-brain samples as well as the large recovery of cells in our experiment. This yielded a total of 48 cell clusters, which were used for cell type-specific soldier-forager differential gene expression testing as well as the regulatory dissimilarity analysis, presented below.

We detected DEGs for each cluster using the MAST algorithm with default settings^45^.We performed two DEG analyses to determine 1) genes that defined each cluster (independent of behavioural state), 2) genes that differentiated soldiers and foragers in each cluster. Cluster-defining genes were used to annotate the 48 detected clusters, and cluster-specific soldier-forager DEGs were used for overlap analyses with colony aggression genes and eGenes.

We annotated many of the 48 detected cell clusters using established insect neuronal and glial biomarkers^25,46,47^. Utilizing available insect scRNA-seq brain atlases of gene expression^18,25,47–49^, including a previous honeybee brain scRNA-Seq analysis^17^, we annotated cell clusters as populations of cells that represented the mushroom bodies (MB), a higher-order multimodal sensory integration center important for learning and memory, navigation, and social behaviour, which formed a clearly segregated set of clusters. Inner-compact Kenyon cells (KCs) were identified by co-expression of *E74* and *ecdysone receptor* while non-compact KCs were identified by co-expression of *Mblk-1* and *dlg5*^50^. Other KC populations detected via expression of *mushroom body-expressed* could not be further annotated, which is expected considering limited knowledge of cellular diversity in the MB. We could also identify cells in the lateral (*ventral veins lacking*) and antennodorsal (*abnormal chemosensory jump 6*) projection neurons of the antennal lobe ^49^. Numerous cell populations constituting the optic lobe were also identified and annotated at superficial levels, as more sequencing will be necessary to further identify all the subpopulations of this extremely heterogenous brain region. Still, we identified populations that were specifically enriched for *Lim3, dorsal root ganglia homeobox*, and co-expression of *homeobox protein homothorax* and *brain-specific homeobox*, markers that span the lobula, medulla, and lamina of the optic lobe^25^. We also found that roughly 88% of cells expressed the neuronal marker *fne* while an average of ∼11% of cells expressed insect glial markers *repo* and *borderless*, a neuron:glia ratio consistent with previous reports from other insects ^18,25^. Within the glial populations, we identified most known insect glia subtypes^51^ including astrocyte-like glia, perineurial glia, subperineurial glia, and cortex glia, as well as a small population of hemocytes (*hml*) that clustered close to the glial populations^25,52^.

### Gene regulatory network analyses: regulatory dissimilarity as a function of variation in colony aggression

To infer single-cell gene regulatory networks (GRNs), we used SimiC, which has been previously shown to perform well with honeybee brain scRNA-Seq data and is better optimized for comparing GRNs across behavioural states than other established methods, to which it has been benchmarked^15^. SimiC imputes scRNA-Seq expression data to infer the relationship between a transcription factor (TF) and its predicted TGs. As input, we used a list of 193 honeybee TFs, each with an robust one-to-one ortholog to a biochemically or genetically validated TF in *D. melanogaster*, which was generated by intersecting TFs predicted by ASTRIX in three previous honeybee brain transcriptomics studies^20,32,53^. We then used SimiC to output a set of incidence matrices containing TF-by-TG (TF x TG) information representative of soldier and forager behavioural states. The GRN evaluation pipeline was based on a weighted area under the curve (“wAUC”) score. For each cell, the wAUC score quantified the relative activity of a given regulon (i.e. a row of the GRN incidence matrix, inferred in the previous step) with respect to the expression of all TGs. For any given TF, if the overall expression of its associated TGs in a cell was high and weights between the TF and its TGs were large, then we concluded that the TF in question has a large influence on the expression profile of a particular cell. For all clusters of cells and TFs, the wAUC was computed independently for each cell yielding a distribution of cell-specific wAUC values corresponding to a behavioural state (soldier or forager). To measure the separation of distributions, we calculated the Kolmogorov-Smirnov (KS) distance between distributions of wAUC values for each cell within a cluster. This yielded one value for each TF per cluster, allowing us to describe “dissimilarities” in soldier and forager regulatory relationships for each predicted cell cluster. Finally, for each cluster, we aggregated the KS distance of each TF into one unique value by computing the area under the curve of the empirical cumulative distribution function (CDF). This approach allowed us to make general comparisons of all cell clusters sampled from soldiers and foragers as well as examine GRN differences specific to the aggression level of the colony of origin.

### Gene regulatory network analyses: architecture of subnetworks associated with colony aggression genes

To infer regulatory subnetworks, we applied the GENIE3^54^ pipeline to both our bulk and sc datasets, using the same list of 193 TFs as the previous step. For soldiers and foragers, we inferred separate GRNs by collecting the top ***N*** regulators (sorted by GENIE3) having a “≥ **C**_**1**_” absolute Spearman correlation with the TG’s expression. Next, we compared soldier and forager GRNs to obtain interactions that are present in both groups with the same TF-TG correlation directionality; this reduced the likelihood of identifying variation due to under-sampling in either soldier or forager groups. We repeated this analysis for bulk and sc expression data. For the sc analysis, we first imputed the expression matrix using MAGIC^55^ with diffusion (“t”) = 2 and split the results into 26 matrices, one for each cell cluster (clustering resolution was set 0.2 in Seurat, with all other parameters unchanged, in order to obtain enough cells per cluster for the analysis). For bulk data, we used ***N*** = 10, **C**_**1**_ = 0.5, and for sc analysis we used ***N*** = 20, **C**_**1**_ = 0.25. Next, we specifically extracted TF-TG relationships in which TGs were either the 30 eGenes (described above) or 15 permutations of 30 random eQTL-genes that were also identified in the eQTL analysis but were not the 30 eGenes (control).

We then assessed similarity of GRN architecture across cell types in eGene and control subnetworks. First, we extracted from each cell cluster a binary vector in which each element corresponded to a TF-TG interaction and compared these vectors across all cell clusters, using Jaccard index as a measure of closeness. Matrices containing Jaccard indices from pairwise comparisons made across cell clusters for eGene or control subnetworks in soldiers and foragers were hierarchically clustered using complete-linkage clustering of the Euclidean distance, and dendrograms for the eGene and control subnetworks were cut at a height of 1.2. After cutting, we specifically explored connectivity of “blocks” (i.e. groups containing more than three cell clusters sharing similar GRN subnetwork architecture based on the above parameters); we identified three blocks from the eGene subnetwork and no blocks from the control subnetwork.

The strength of connectivity (i.e. similarity of subnetwork architecture across cell clusters) was assessed by assigning a score to each block in the eGene subnetwork representing the smallest pairwise connectivity score observed between clusters within a block. This score was then computed in each of the 15 permutation subnetworks using the same cell clusters identified in each respective block in the eGene subnetwork. The minimum connectivity scores identified in the eGene subnetworks were considerably higher than scores generated in each of the permutation networks.

TF rank was computed by dividing each TF’s appearance in a given subnetwork by the total number of interactions in the subnetwork. TF-TG correlations that differed ≥ 0.1 between soldiers and foragers represented ∼10% of all interactions in the eGene subnetwork and were therefore considered to be the strongest differences.

**Fig. S1.**
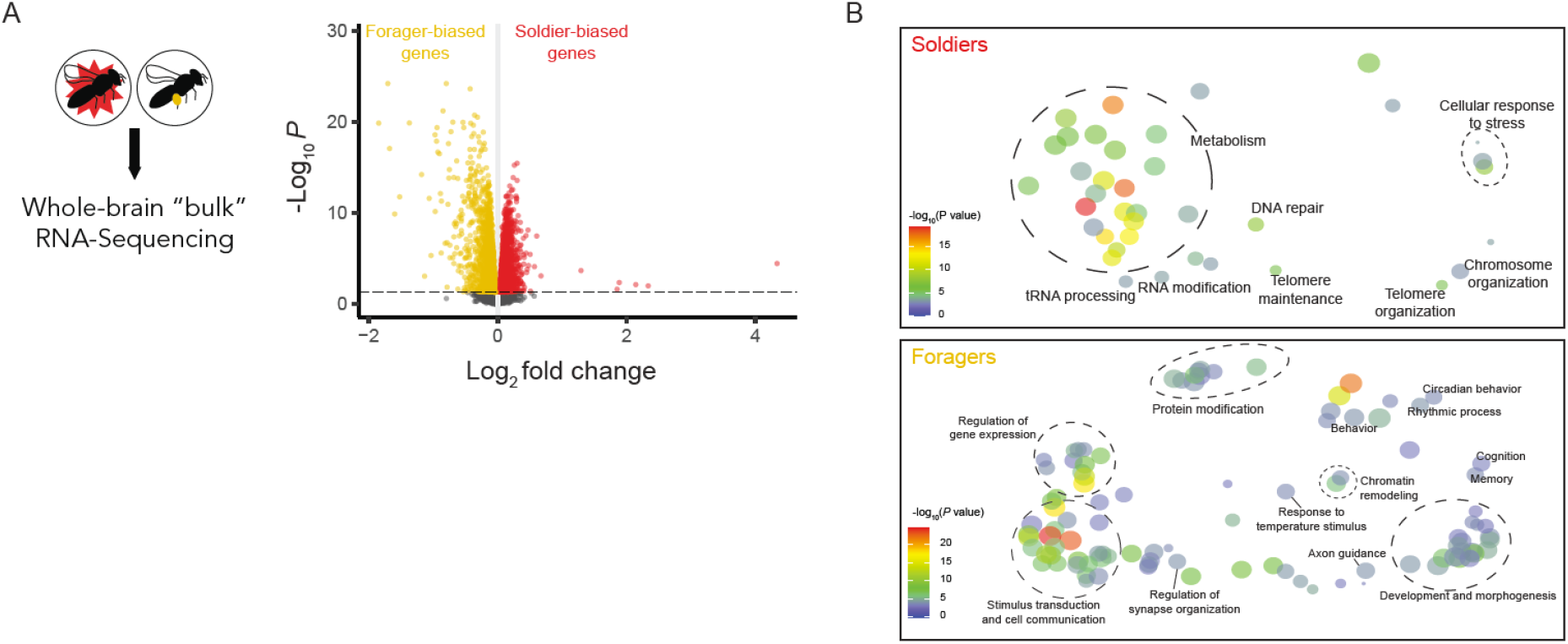
Whole-tissue brain transcriptomics of soldiers and foragers. (A) Whole-brain RNA-Sequencing of the same soldier and forager bees from ^2^ revealed thousands of differentially expressed genes (DEGs) associated with behavioural state, termed “soldier-” or “forager-biased” based on fold-change following a pairwise comparison between groups. Red or yellow dots represent genes upregulated in soldiers or foragers, respectively, that passed a false discovery rate (FDR)-corrected *P* value threshold of 0.05; grey dots represent tested genes that did not pass this threshold. A total of 4278 DEGs were identified, with 2242 and 2036 that were soldier-or forager-biased, respectively (Data S1). (B) REVIGO plots show Biological Process terms identified by a Gene Ontology (GO) enrichment analysis associated with soldier-or forager-biased gene lists. Circles are organized in semantic space such that more similar terms in the GO hierarchy are positioned more closely together. Circle diameter is inversely correlated with the specificity of each GO term, such that smaller circles represent more specific terms.

**Fig. S2.**
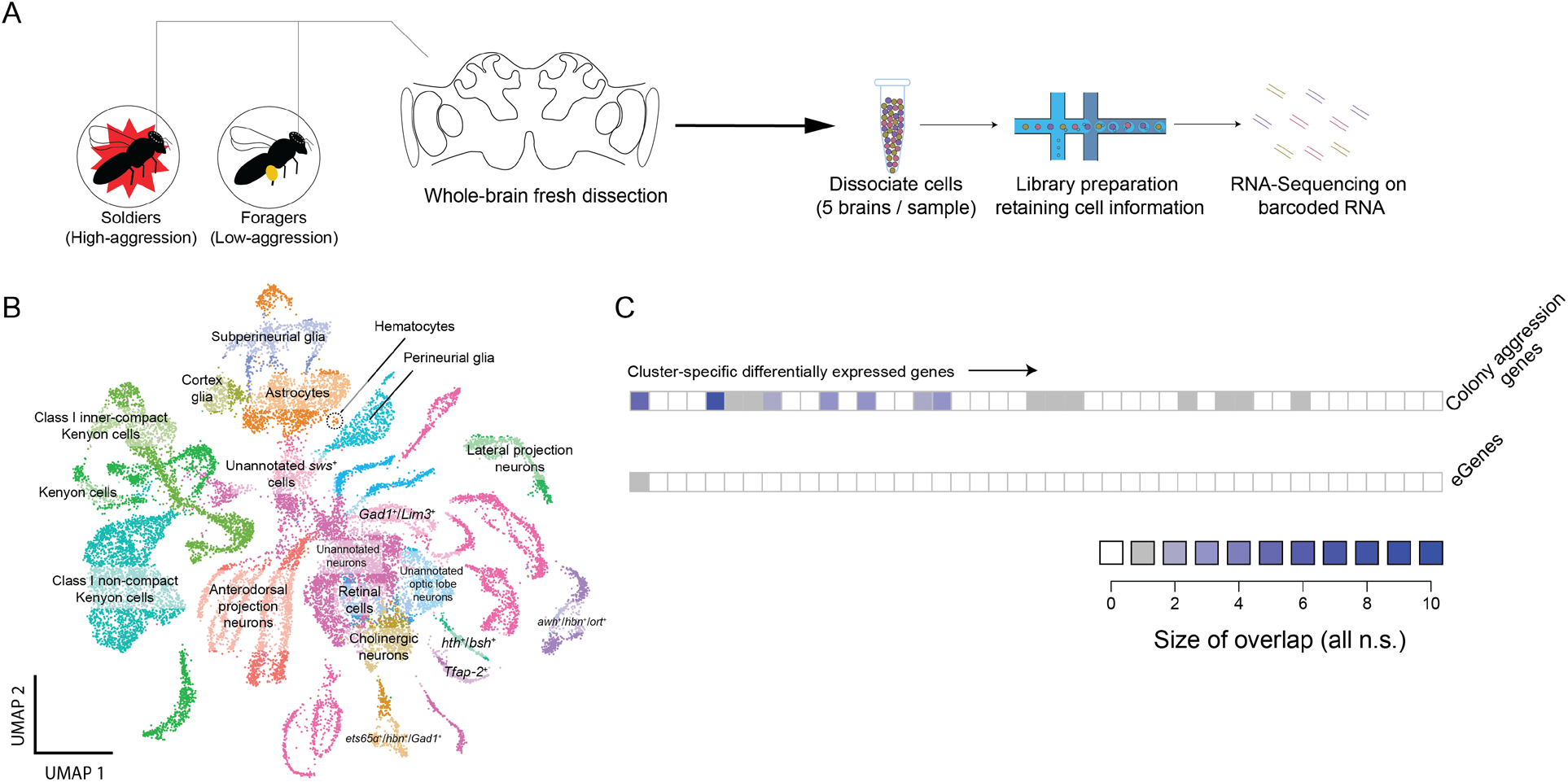
Single-cell brain transcriptomics of honeybee soldiers and foragers. (A) Workflow for transcriptomic profiling of ∼24,000 cells from the brains of field-collected soldiers and foragers and (B) bioinformatic clustering of cells in reduced dimensionality (UMAP) space. Clusters were manually annotated using cell type-specific markers as described in Methods. (C) Cluster-specific differentially expressed genes between soldiers and foragers did not significantly overlap with colony aggression genes from ^2^ or eGenes (hypergeometric test, FDR > 0.10 for all comparisons). Color represents size of overlap for each cluster; clusters with no soldier-forager DEGs not shown.

**Fig. S3.**
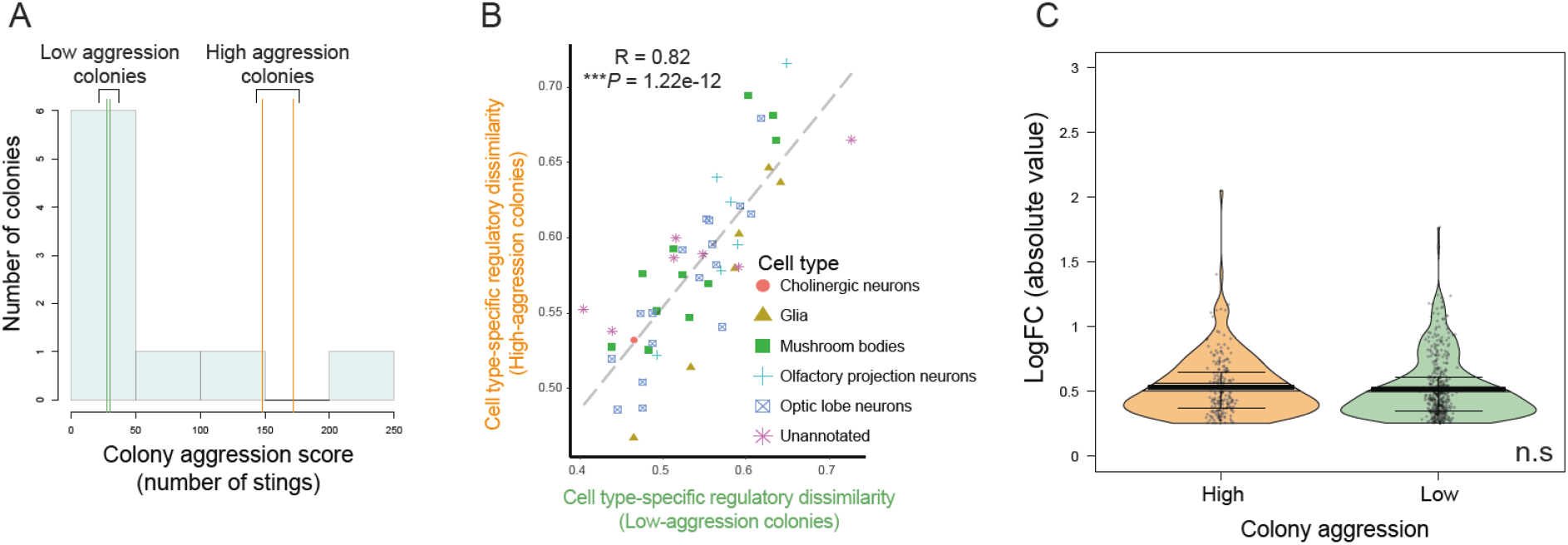
Single-cell RNA-Sequencing reveals that regulatory dissimilarity, but not differential expression, changes across soldiers and foragers as a function of colony aggression. (A) We divided our scRNA-Seq comparisons of soldiers and foragers by colony aggression level, considering colonies in the top 15% or bottom 25% of the aggression distribution reported in ^2^ to be “high-” (orange vertical lines) and “low-aggression” (green vertical lines) colonies, respectively, with soldier and forager samples from two colonies in each aggression category. X-axis refers to number of stings delivered by resident aggressors to small leather patches placed in front of the entrance during colony disturbance (Methods). (B) The rank-order of cell type-specific regulatory dissimilarity (RD) scores, as described in Fig. 1E, was preserved across high-and low-aggression colonies (Pearson’s product-moment correlation). (C) Unlike for RD, as shown in Fig. 2A, there was no difference in the fold-change magnitude of cell type-specific differentially expressed genes between soldiers and foragers from high-and low-aggression colonies (Wilcoxon rank-sum test with continuity correction, *P* > 0.10); gene regulatory relationships, rather than expression alone, may therefore be more appropriate for linking group and individual at the molecular level.

**Fig. S4.**
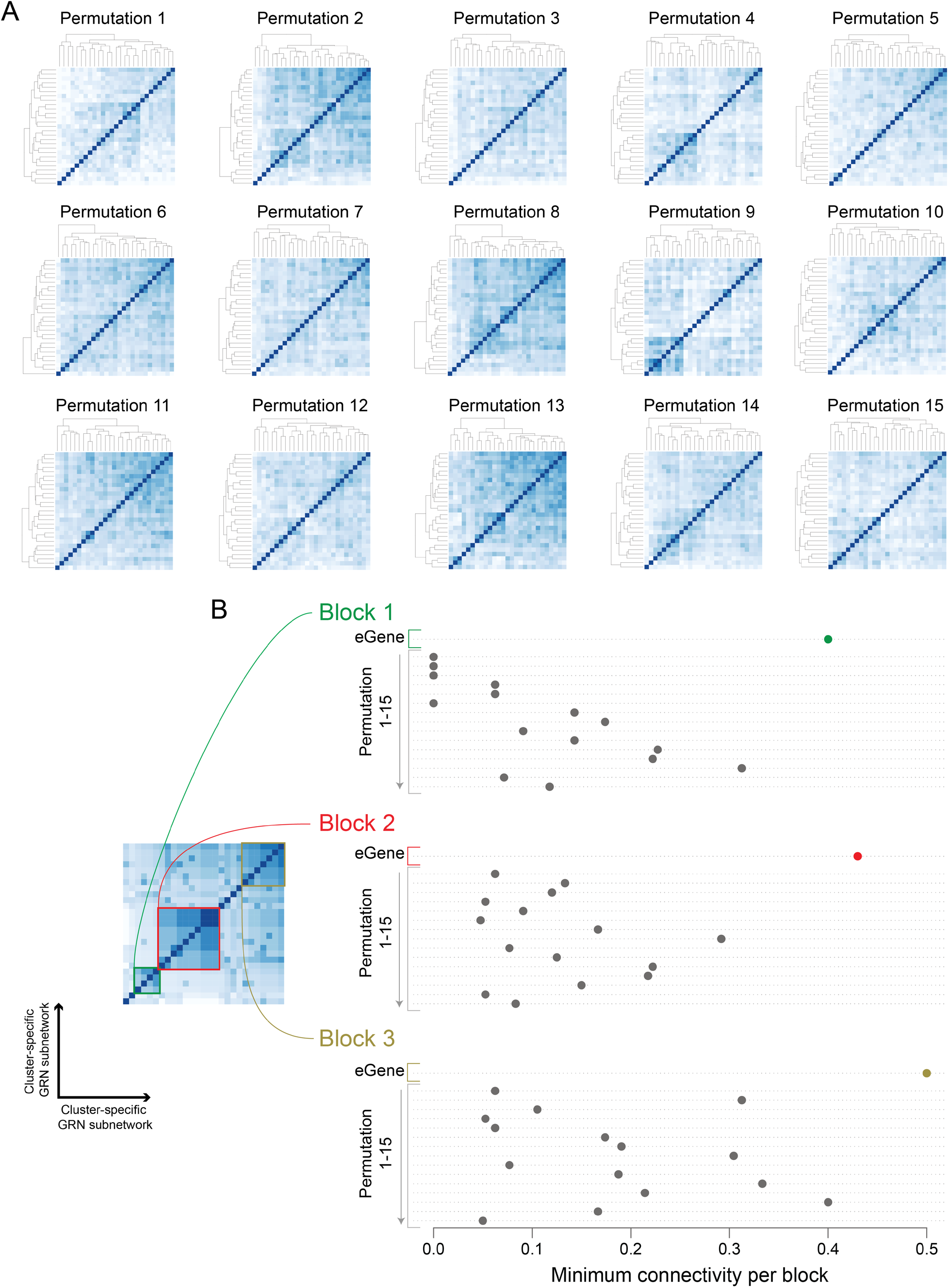
Random permutation subnetworks are less connected than eGene subnetwork. (A). Heatmaps as described in Fig. 2 for each of the 15 random permutation subnetworks. (B). Using the three “blocks” of sc clusters identified in Fig. 2, dot charts compare smallest (“minimum”) pairwise connectivity for eGene or 15 control subnetworks. For each block, the minimum connectivity was much higher in the eGene subnetwork compared to the 15 controls.

**Fig. S5.**
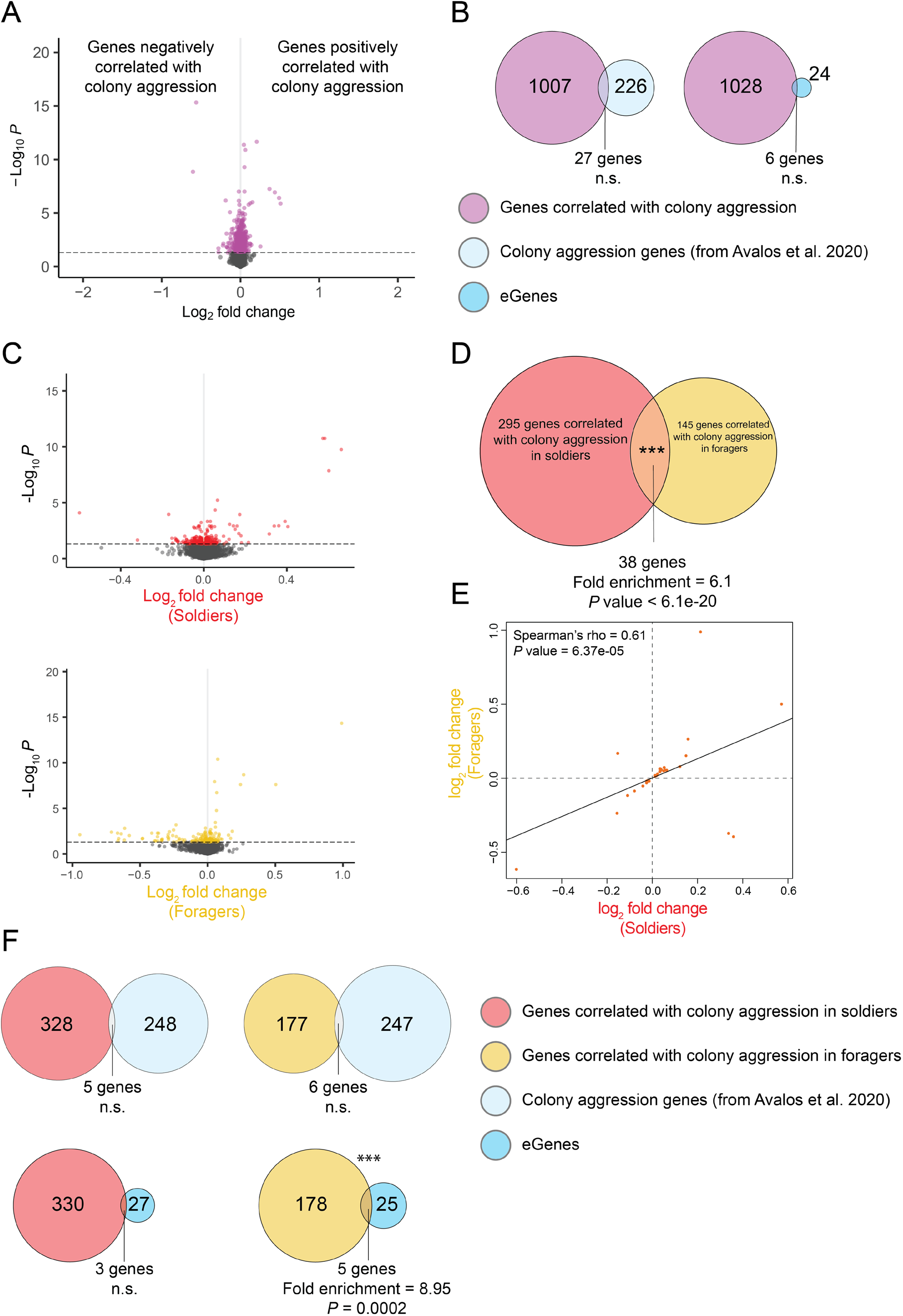
Molecular correlates of colony aggression identified in soldier and forager whole-tissue brain transcriptomics. (A) We regressed brain gene expression on colony-level aggression for each individual soldier and forager in a given colony, identifying 1037 genes correlated with colony aggression. (B) None of these genes were found to significantly overlap with colony aggression genes from ^2^ or eGenes identified in this study. (C) Going deeper, we performed the same analysis independently in soldiers and foragers (fold-change on volcano plots as described in [A]) and (D) identified a significantly overlapping set of genes that were (E) highly concordant in terms of differential expression associated with colony aggression. (F) Performing an overlap analysis now resolved to the level of soldiers and foragers identified a significant overlap between genes correlated with colony aggression in foragers and eGenes; no other comparison was significant (hypergeometric test, *P* > 0.1).

